# Stochastic modeling of aging cells reveals how damage accumulation, repair, and cell-division asymmetry affect clonal senescence and population fitness

**DOI:** 10.1101/628669

**Authors:** Ruijie Song, Murat Acar

**Author notes:** **E-mail addresses of the authors:** Ruijie Song, Murat Acar. **Full postal address of the submitting author:** Murat Acar, Systems Biology Institute, Yale University, 850 West Campus Drive, Room: ISTC-122, West Haven, CT 06516, U.S.A.

## Abstract

**Background:** Asymmetry during cellular division, both in the uneven partitioning of damaged cellular components and of cell volume, is a cell biological phenomenon experienced by many unicellular organisms. Previous work based on a deterministic model claimed that such asymmetry in the partitioning of cell volume and of aging-associated damage confers a fitness benefit in avoiding clonal senescence, primarily by diversifying the cellular population. However, clonal populations of unicellular organisms are already naturally diversified due to the inherent stochasticity of biological processes.

**Results:** Applying a model of aging cells that accounts for natural cell-to-cell variations across a broad range of parameter values, here we show that the parameters directly controlling the accumulation and repair of damage are the most important factors affecting fitness and clonal senescence, while the effects of both segregation of damaged components and division asymmetry are frequently minimal and generally context-dependent.

**Conclusions:** We conclude that damage segregation and division asymmetry, perhaps counterintuitively, are not necessarily beneficial from an evolutionary perspective.

## BACKGROUND

Even though the somatic cells of multicellular organisms accumulate significant amounts of aging-related damage throughout the lifetime of the organism, their young progeny generally start with low levels of protein damage. A number of mechanisms have been proposed to explain this phenomenon, generally involving some special way of eliminating damage in germ cells, or elimination of damaged germ cells [1–5]. A major hallmark of aging in higher organisms is the depletion or dysfunction of stem cells, which accumulate various forms of molecular damage during the aging process [6–8]. For unicellular organisms such as the budding yeast *Saccharomyces cerevisiae* or the fission yeast *Schizosaccharomyces pombe* undergoing mitosis, there is no somatic/germ cell distinction. Yet both *S. cerevisiae* and *S. pombe* exhibit lineage-specific aging [9–14]. In the budding yeast, for instance, the mother cell is known to accumulate aging-related damage markers such as extrachromosomal rDNA circles (ERCs) as it ages and eventually enters replicative senescence and dies, while the daughter cells that bud off from the mothers are mostly protected from the accumulated ERCs and generally enjoy a full replicative lifespan even if born from old mother cells [15]. Similar segregation of damaged proteins has been observed during the binary fission of *S. pombe*, where oxidatively damaged carbonylated proteins are concentrated in one of the two daughter cells – in this case, the one carrying the previous birth scar [16]. Lineage-specific aging has also been observed in the bacteria *Caulobacter crescentus* and *Escherichia coli* [10, 17–20], leading some to analogize the lineage-specific aging behavior in unicellular organisms to the somatic/germ cell distinction in higher ones.

The observation of damage-partitioning behavior even in unicellular species naturally raises the question of whether there is any selective advantage resulting from it. Using a deterministic model based on ordinary differential equations (ODEs) with fixed parameter values, Erjavec and colleagues examined two forms of asymmetry during the cell-division process, which we will denote as *division asymmetry* and *damage segregation,* respectively. The former refers to the asymmetric partitioning of cell volume (as naturally seen in *S. cerevisiae*) between the two cells after division, while the latter refers to the asymmetric partition of damaged cellular components. They concluded that such behavior indeed confers a fitness advantage: under their model, both damage segregation and division asymmetry allowed the population to survive a higher level of protein damage without entering clonal senescence than it otherwise would have [16]. The researchers attributed this effect to the ability of these mechanisms to diversify individuals within a population; in the absence of these mechanisms, the population of cells in their simulations are entirely homogeneous [16].

The fact that real cells are not homogeneous at all raises questions about the reliability of these predictions made based on such a fully deterministic model. It is well known that the expression level of genes can fluctuate substantially, even among cells that are genetically identical or indeed in the same cell over time [21–27]. This kind of fluctuations, commonly known as noise, can come from a variety of sources: cell-to-cell variations in the abundance of transcription and translation machinery (such as RNA polymerase, general transcription factors, and ribosomes), for instance, or the stochastic nature of transcription events that take place in any single cell [21, 28]. Indeed, it has been shown that stochastic noise can cause drastic differences between reality and what a deterministic model predicts [29].

Bringing a more realistic approach to the study of aging cells in terms of damage accumulation, segregation behavior, and their effects on clonal senescence and fitness, here we investigate whether, and under what circumstances, damage segregation and division asymmetry confer fitness advantages in freely dividing unicellular organism populations when noise is taken into account. We focus on two forms of fitness advantages: resistance against clonal senescence, and increased rates of population growth. We find that damage mitigation and the rate of damage accumulation play major roles in determining the fitness of the cells.

## MATERIALS and METHODS

### Modeling of cell growth and division in aging cells

We consider a cell that grows exponentially in volume during the cell cycle [30] and accumulates damage as it grows (Fig. 1A). Cells are assumed to accumulate damage (*D*) at a constant rate *r*_*dmg*_, and reduce damage via two sources, actively by repair and passively by dilution due to cell division and volume growth:

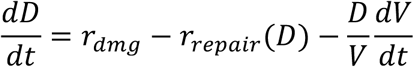

**Fig 1.**
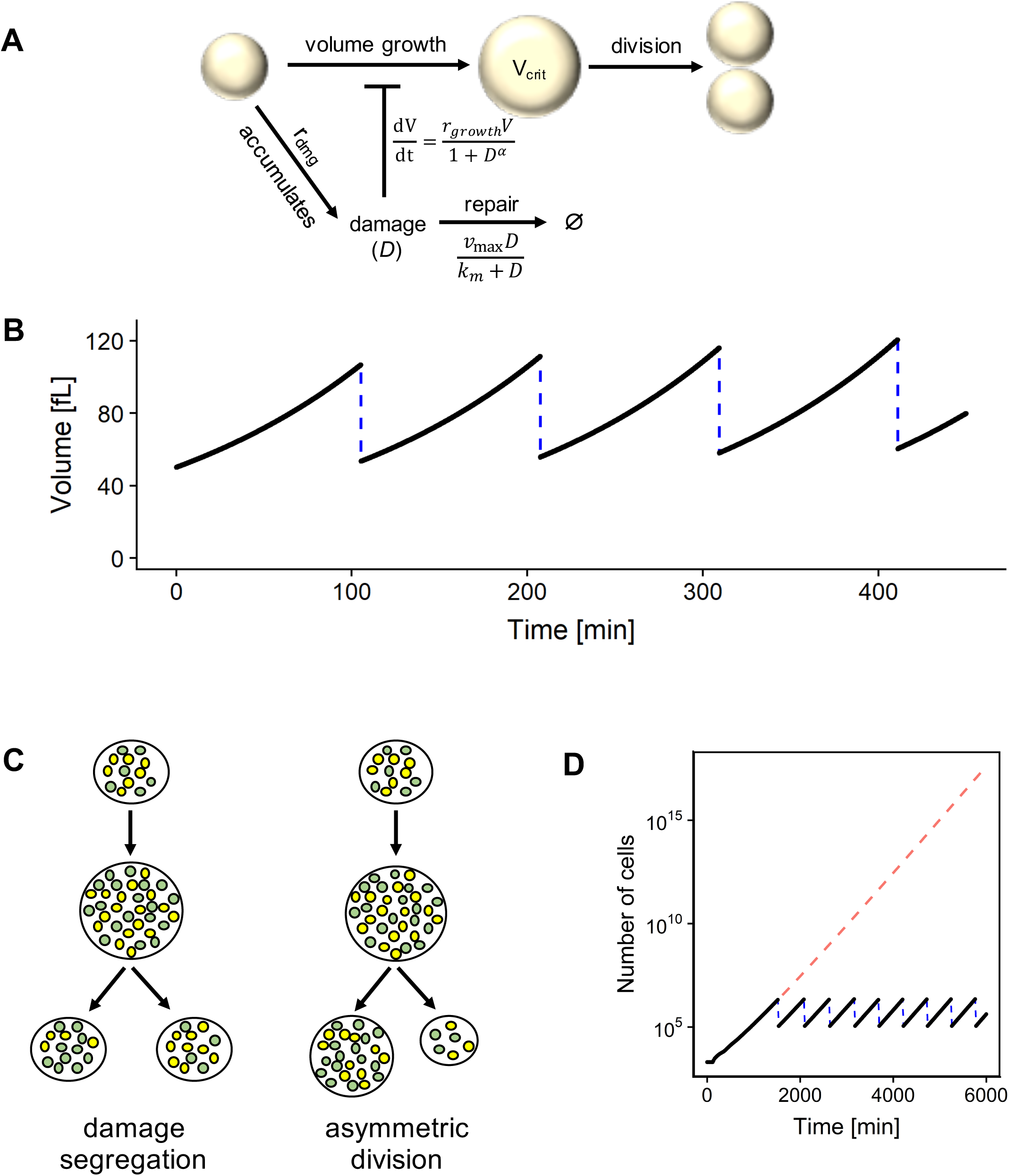
**A.** Illustration of the model. The cell grows until it reaches a critical volume and divides. It accumulates damage over time, which slows down volume growth. The accumulated damage can also be repaired. There is no separate mechanism for killing a cell due to damage, because high level of accumulated damage will prevent a cell from dividing and cause it to be rapidly overtaken by faster-dividing cells. **B.** The cell volume module of the model. The volume grows exponentially until it reaches a generation-dependent critical volume and the cell divides (blue dashed line). **C.** Two mechanisms of particular interest in this study: segregation of damaged proteins in mother cells, and division asymmetry of cell volume. Yellow dots indicate normal proteins, while green dots indicate damaged proteins. **D.** Illustration of the exponential growth of simulated cell population. An initial population of 2000 cells were simulated for 6000 minutes with periodic resampling (blue dashes) every time the population size exceeds 2 million cells. The red dashed line indicates the expected population size without sampling.

Here, *D* is an abstract value representing the concentration of damaged cellular components and other harmful artifacts of aging. For simplicity, we assume that the forms of damage represented by *D* are freely diffusible, segregable, and repairable, and that there are no other sources of damage affecting cell growth. The rate of damage repair *r*_*repair*_ as a function of *D* is assumed to follow Michaelis-Menten kinetics with parameters *v*_max_ and *k*_*m*_:

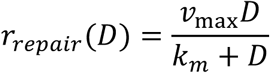

The cell volume (*V*) grows exponentially at a rate that is slowed by accumulated damage:

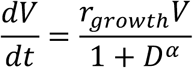

where *r*_*growth*_ is the maximum growth rate constant and *α* is a nonlinearity coefficient.

In the initial population of cells, each cell starts at an initial volume *V*_*i*_. A cell is assumed to divide when it reaches a generation-dependent critical volume *V*_*crit*_ (Fig. 1B). This critical volume increases linearly with replicative age (Table S1), consistent with the observations on single budding yeast cells [27]. The parameters of volume growth during cell cycle were selected to roughly correspond with the microscopic growth dynamics measured in budding yeast cells (Table S1) [31], with an approximate expected damage-free doubling time of 100 minutes for symmetrically dividing cells. We separated global noise into two categories: noise in cell volume control (*n*_*v*_), and noise in damage and its repair (*n*_*d*_). In each case, global noise was simulated as a random perturbation applied to each corresponding parameter: the initial parameter value of each individual cell was sampled from a normal distribution *N*(*μ* = *p, σ* = *np*), where *p* is the selected mean parameter value from Tables S1-S2 and *n* is the applicable noise level.

During cell division, we consider the original cell (“mother”) to retain its identity and produce a new daughter cell, for ease of reference. The accumulated damage is distributed between mother and daughter cells as follows. Let *D* be the damage level of the mother cell before division, then the damage level of the newly produced daughter cell is equal to (1 – *s*)*D*, where *s* in the range [0, 1] is the parameter quantifying the extent of damage segregation, and the damage level of the mother cell after division is 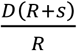, where *R* is the ratio of the volume between the mother and daughter components after division. In other words, the total amount of the damage (equal to the damage level times volume) is constant across the specific cell division event.

At each cell division, we assumed that each daughter cell partially inherits its mother’s parameter values for all parameters listed in Tables S1-S2. For each parameter *p*, the new value *p*_*new*_ follows the relationship *p*_*new*_= *c p*_*mother*_+ 1 – *c*) *p*_*fresh*_, where *p*_*fresh*_ is a parameter value freshly sampled from the normal distribution applicable to that parameter, *p*_*mother*_ is the parameter value for the mother cell, and *c* is a constant in the range [0, 1] characterizing the level of inheritance: when *c* = 0, the daughter gets a new parameter value from the same distribution used to generate the parameter values used for the initial population of cells, while when *c* = 1, the daughter perfectly inherits its mother’s parameter value.

Since the simulated cells, just like real ones grown without nutrient limitations, exhibit exponential growth and can easily overwhelm the computing capacity if left unchecked (Fig. 1D), we kept the population size low by means of periodic resampling as a mimicry of using a turbidostat [25]. Because cells slow to divide due to damage are expected to be rapidly overtaken by faster-dividing cells, we did not include a separate procedure for killing cells due to accumulated damage.

To keep the generality of the model intact, we determined the parameter values to use in our model by combinatorially selecting from a grid spanning a wide range (Table S2), with a total of 7.185 million sets of parameter values tested. The parameter bounds are hand-selected to cover the arguably biologically plausible range. The noise parameters were capped at 10% because each parameter is perturbed independently, and so the noise in each parameter is expected to combine to produce higher noise in the overall phenotype. We chose the range of *r*_*dmg*_ so that the maximum will virtually immediately block cell growth in the absence of repair, and then chose the range of the repair parameters to match the range of *r*_*dmg*_. These parameters are also sampled on a logarithmic scale so as to capture a wide variety of damage strengths.

For each parameter set, we recorded its population size trajectory over the course of the simulation. From these numbers we calculated the average doubling time of the population. If the calculated average doubling time was greater than 1500 minutes, it was treated as 1500 minutes for the purpose of analysis. Each simulation was repeated three times and the average doubling time resulting from the three repeats was calculated. For parameter sets producing reasonable fitness levels (<400 min doubling time), we do not observe significant changes in the computed doubling time if the initial 600 minutes of the trajectory is omitted. This is expected since these populations relax rapidly to the steady state and the initial conditions have very limited impact when averaged over the long course of the simulation.

### Simulations of the Stochastic Model

The model as described above was implemented in C++ using custom-written code, utilizing the SUNDIALS library [39]. The complete set of model parameters are summarized in Tables S1-S2.

Simulations for each parameter set chosen according to the tables above were performed from an initial set of 2000 cells. Every 40 minutes, the number of cells in the population was recorded and the fold-change in population growth from the previous time point was calculated, then the population was randomly resampled down to 2000 cells.

For the illustration of exponential growth as shown in Figure 1E, the simulation was performed as described above, except that

- The population size was recorded every 20 minutes;
- Resampling was not performed until the population size reached 1000 times the initial sample size (i.e., 2 million cells);
- The population size for the resampling was 50 times the initial sample size (100,000 cells).

### Analysis of Simulation Results

We quantified the effect of changing the value of a parameter *P* on fitness (resistance against clonal senescence or increased rate of population growth) as follows. For simplicity, we denote the nine parameters of the model *P*_1_, …, *P*_9_, which can take *N*_1_, …, *N*_9_ possible values, respectively (Table S2). Without loss of generality, let *P*_1_ be the parameter *P* we want to examine. Then, we partition the 7.185 million combinations into 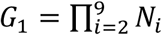 groups of *N*_1_ combinations each, where the combinations in each group only differ in the value of *P*_1_ (and have the same values of *P*_2_, …, *P*_9_). For each group, we then computed a minimum and a maximum doubling time, from which we determined whether changing the value of *P*_1_ for this particular set of parameter value combination could cause a significant change in fitness (for clonal senescence, a difference in outcome; for growth rate, defined as more than 5% difference between minimum and maximum doubling time). Repeating this for all *G*_1_ groups, we found that, in *M*_1_ of them, changing the value of *P*_1_ resulted in a significant change in fitness. Then, the frequency at which a change in the value of *P*_1_ could significantly alter fitness was *M*_1_/*G*_1_. The total number of groups for all parameters combined is *G* = Σ *G*_*i*_. = 12.56 million. For clonal senescence, we found changes in *M* = Σ *M*_*i*_ = 1.41 million groups. For fitness, we found significant changes in *M* = Σ *M*_*i*_ = 2.38 million groups.

### Quantification of Relative Abundance

Each panel in Figures 3-7 and Figures S1-S4 quantifies the relative abundance of each possible value of a parameter *Q* among the parameter combinations where changes in the value of a different parameter *P* has a significant effect (>5%) on fitness. We denote the possible values of *Q* as *Q*_1_, …, *Q*_*n*_, and also denote the number of combinations where *Q* = *Q*_*i*_. and changes in *P* can cause a change (>5%) in fitness as 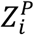. We further partition 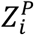 into three groups (colored blue, green and red in the figure panels) based on the value of *P* at which maximum fitness is attained (i.e., 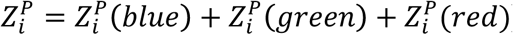), blue if *P* is at its largest possible value, red if *P* is at its smallest possible value, and green if *P* is at an intermediate value.

**Fig 2.**
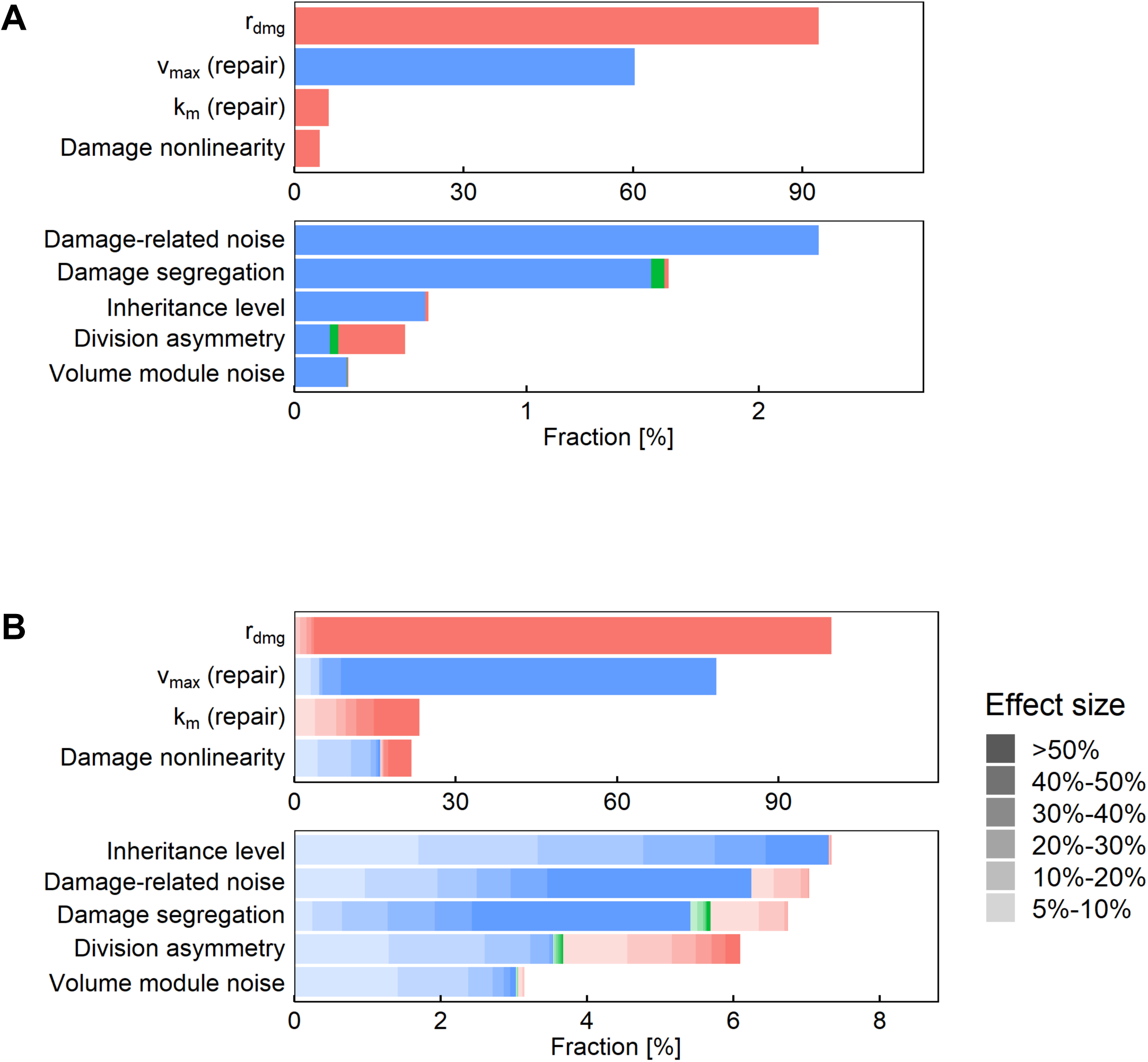
**A.** Sensitivity analysis for the clonal senescence outcome. The parameter combinations tested were partitioned into groups such that all combinations in each group differs only in the value of one parameter. The fraction of groups with divergent senescence outcome is plotted for each parameter. **B.** Sensitivity analysis for the doubling time outcome. The parameter combinations tested were partitioned into groups such that all combinations in each group differs only in the value of one parameter. The fraction of groups where the minimum doubling time is at least 5% below the maximum is plotted for each parameter. Level of transparency indicates the size of the effect. In each panel, the color indicates the value of the parameter corresponding to the most-fit combination in the group: red indicates that the smallest parameter value is the most fit; blue indicates that the largest parameter value is the most fit; and green indicates that an intermediate parameter value is the most fit.

**Fig 3.**
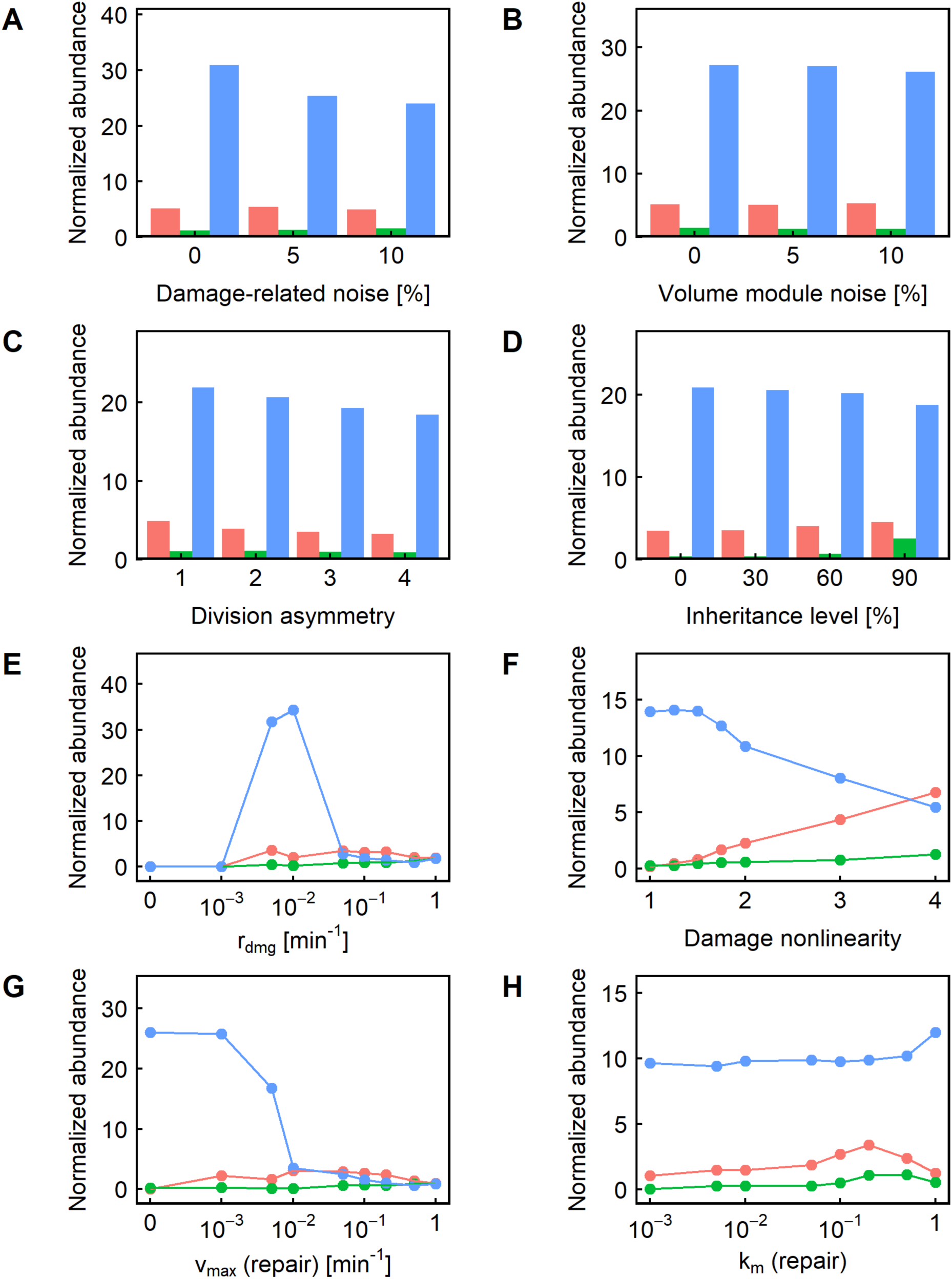
**A-H.** Relative representation of the other parameters in the cases where changing the level of damage segregation caused a significant (>5%) fitness difference. In each panel, the color indicates the level of damage segregation resulting in maximum fitness: red indicates that no segregation is the most fit; blue indicates that full segregation is the most fit; and green indicates that an intermediate level of segregation is the most fit.

Since the possible values of *Q* are arbitrarily selected from a large grid, the values are not necessarily equally responsive to fitness changes. Thus, as a normalization measure, we normalized the value of 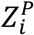 (and the partitioned) by the total number of combinations where *Q* = *Q*_*i*_. and changes in the value of any other parameter can cause a change (>5%) in fitness. In other words, the normalized value is 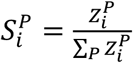. Similarly, for each color *C* the normalized value is 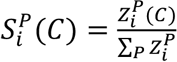.

The value of 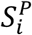 can vary significantly depending on the identity of the parameter *P*. To make the abundance value more uniform across panels, we further multiplied 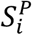 and 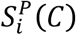 for each color by a scaling factor equal to 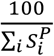 to produce the normalized abundance of *Q*_*i*_. plotted in each panel. Thus, the normalized abundance points within in each panel add up to a constant value of 100.

## RESULTS

### Dissecting the key parameters affecting the clonal senescence outcome

In our model, a cell reaches *senescence* when its growth rate is slow enough as to virtually stop dividing. A clonal population of cells exhibits *clonal senescence* if every single cell in the population reach senescence. For the purposes of our analysis, we classified a cell population as clonally senescent if it exhibits an average doubling time greater than 1000 minutes over the course of the simulation, which is more than ten times the expected doubling time in damage-free conditions.

To determine the degree of importance of a model parameter for clonal senescence, we examined how likely it is for changes in the value of one parameter to alter the senescence outcome. More formally, for each parameter *P*, we partitioned the 7.185 million parameter value combinations into disjoint groups such that the combinations in each group only differ in the value of *P*, and calculated the fraction of groups whose combinations diverge in the clonal senescence outcome. While the absolute value of this fraction is necessarily dependent on the values of the other parameters we picked in the study, the relative value is still indicative of whether the parameter is relevant generally, or only when the other parameters are in a relatively narrow region of the parameter space.

Changes in the damage rate *r*_*dmg*_ caused a different senescence outcome in 93% of parameter combinations, and changes in the repair rate *v*_max_ caused a different outcome in 60% of cases. These observations were intuitive and reaffirmed the validity of the model setup, as the strongest effect was exerted on the amount of accumulated damage, with the possible values of the parameters spanning a wide range. As expected, we find the most-fit combination to be when the damage rate is lowest and the repair rate is highest (Fig. 2A, as indicated by red and blue coloring of their respective bars).

Changes in damage-related noise (*n*_*d*_), the Michaelis constant (*k*_*m*_) for repair, and the nonlinearity of damage’s effect on growth (*α*) are less likely to affect the clonal senescence outcome. In only 4.6% of the parameter combinations did a change in *α* affect the clonal senescence outcome; for *k*_*m*_, the number is slightly higher at 6.1%, while for *n*_*d*_, it is lower at 2.3%. These parameters also affect the amount of accumulated damage or its effect on the cell, but the effects are weaker and less direct. When changes in these parameters did affect the senescence outcome, the direction is essentially uniform: in almost all of the cases, clonal senescence is avoided by having high noise, low *k*_*m*_, or low *α* (Fig. 2A, color). This again makes sense: a lower *k*_*m*_ means a higher repair rate, while a lower *α* means a higher growth rate (when *D* > 1, which is necessary for clonal senescence to even come into play because if *D* < 1 then the volume growth rate can’t fall below half of the maximum growth rate). In the borderline cases where noise matters, moreover, higher noise means a better chance to come across good parameter values that could sustain the population.

Damage segregation and division asymmetry only affected the clonal senescence outcome in a very small fraction of parameter combinations – 1.6% and 0.4% respectively (Fig. 2A). We did find that damage segregation is overwhelmingly beneficial in the few cases where it did matter: in 99% of the cases in which segregation made a difference on the senescence outcome, some damage segregation (represented as the blue and green portions of the bar) was needed to avoid clonal senescence (Fig. 2A). On the other hand, division asymmetry is more likely to be detrimental, if not overwhelmingly so: in 60% of the cases where asymmetry made a difference in the senescence outcome (represented by the red portion of the bar), lack of asymmetry is necessary to avoid clonal senescence, while in the remaining 40% some level of asymmetry is necessary.

We therefore conclude that damage segregation and division asymmetry are not the main effectors of the senescence outcome. Interestingly, neither mechanism is capable of altering the total amount of damage accumulated in the population, which appears to be the key determinant. Thus, changing the damage accumulation rate and the maximum repair rate are most likely to cause (or avoid) clonal senescence.

### Characterizing the effect of age-associated damage on population fitness

Clonal senescence, which implies a complete loss of fitness, is a drastic outcome, and it is certainly conceivable that a parameter might affect population fitness incrementally without causing the entire population to become senescent. We therefore examined the ability of each model parameter to affect the growth rate (or fitness) of the population (Fig. 2B). For this analysis, we calculated the doubling time output of our model using the parameter sets determined as described in the previous section, and examined the cases where the change in the value of one parameter could lead to a significant change (>5%) in doubling time. For each parameter *P*, we again partitioned the 7.185 million parameter value combinations into disjoint groups such that the combinations in each group only differ in the value of *P*, and calculated the fraction of groups whose maximum and minimum doubling time are different by more than 5%.

#### The parameters most likely to affect the senescence outcome are also most likely to have strong fitness effects

As in the output of clonal senescence, we found that *r*_*dmg*_ and *v*_max_ were the two parameters most likely to cause a fitness differential (>5% in terms of doubling time). Changing the damage accumulation rate *r*_*dmg*_ is capable of significantly altering fitness in more than 99% of all parameter combinations used, while changes in the maximum repair rate *v*_max_ significantly altered fitness in 78% of the parameter combinations (Fig. 2B). Other damage-related parameters are also more likely to affect fitness: changes in *k*_*m*_ and *α* are each capable of affecting fitness in about one-fifth (23% and 22%, respectively) of the parameter combinations tested, compared to 3% for the volume-module noise, the parameter that turned out to be the least likely to cause fitness differences (Fig. 2B). For three of the four parameters, moreover, the effect of a parameter value change on fitness is monotonic (as indicated by the prevalence of one color in the figures): a higher *v*_max_ (Fig. S2) virtually always improved fitness, while a higher *r*_*dmg*_ (Fig. S4) or *k*_*m*_ (Fig. S3) decreased fitness. This is expected given the functional forms linking these parameters to the model. *α*, on the other hand, turned out to be a parameter with a double-edged impact on fitness, though unsurprisingly (Fig. 2B, 7, blue and red color). Introducing ultrasensitivity means that some damage levels will have less impact on fitness while others will have more. Depending on the other parameter values, then, the impact of *α* on fitness could and did go in both directions (Fig. 7).

**Fig 4.**
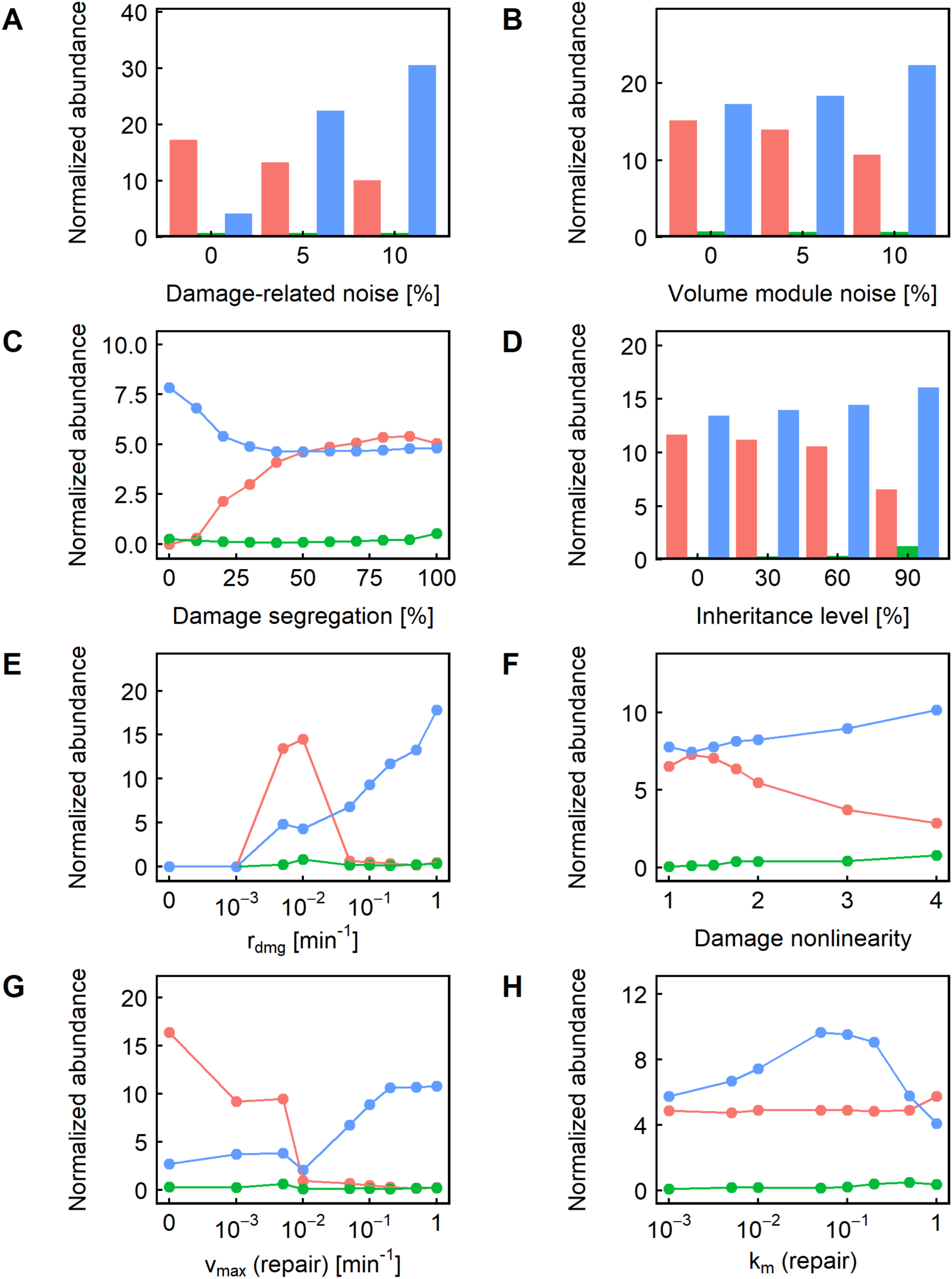
**A-H.** Relative representation of the other parameters in the cases where changing the level of division asymmetry caused a significant (>5%) fitness difference. In each panel, the color indicates the level of division asymmetry resulting in maximum fitness: red indicates that no division asymmetry is the most fit; blue indicates that maximum asymmetry (1:4) is the most fit; and green indicates that an intermediate level of asymmetry is the most fit.

**Fig 5.**
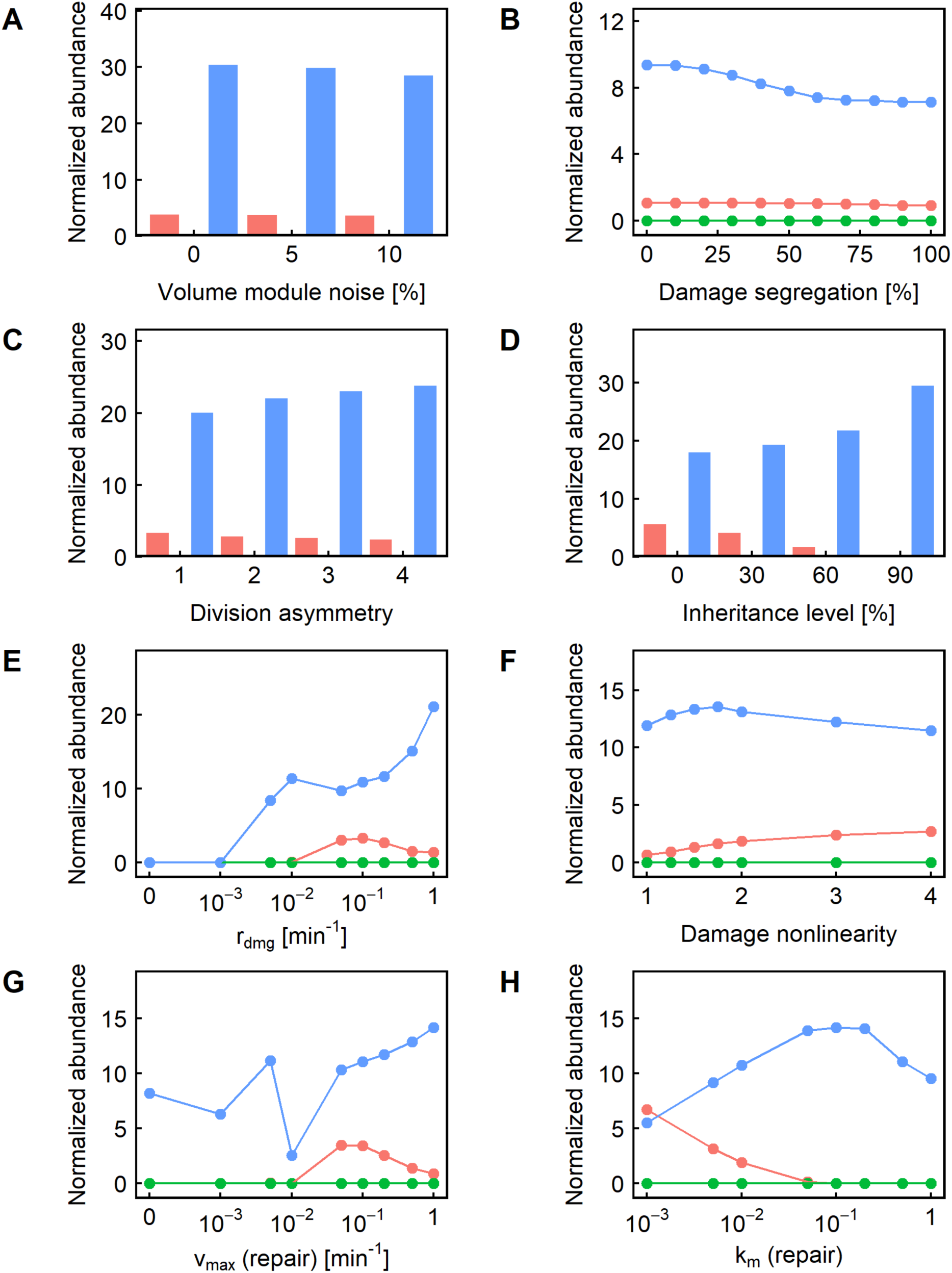
**A-H.** Relative representation of the other parameters in the cases where changing the level of damage-related noise caused a significant (>5%) fitness difference. In each panel, the color indicates the level of damage-related noise resulting in maximum fitness: red indicates that no damage-related noise is the most fit; blue indicates that maximum damage-related noise (10%) is the most fit; and green indicates that an intermediate level of damage-related noise (5%) is the most fit.

**Fig 6.**
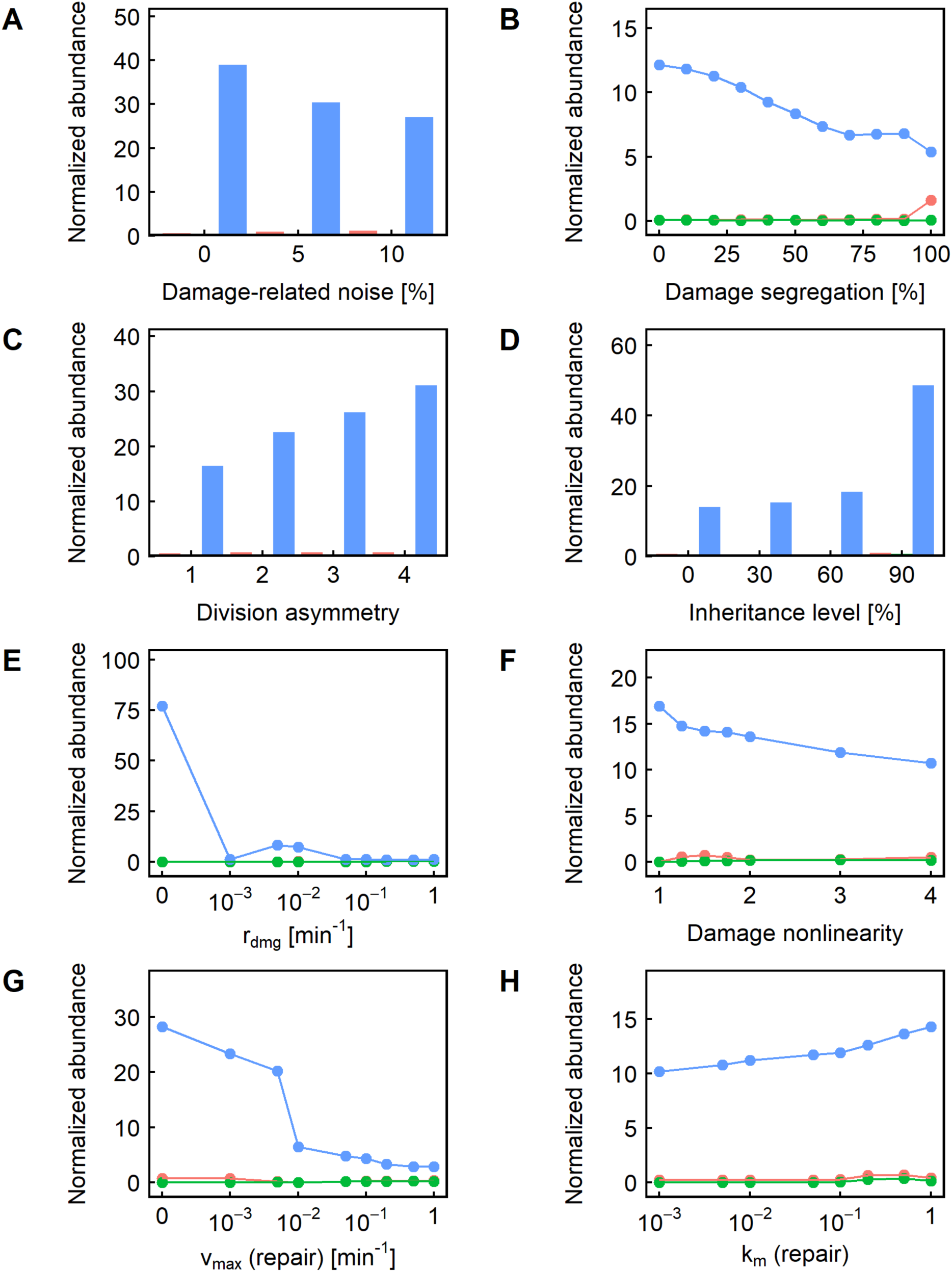
**A-H.** Relative representation of the other parameters in the cases where changing the level of volume module noise caused a significant (>5%) fitness difference. In each panel, the color indicates the level of volume module noise resulting in maximum fitness: red indicates that no volume module noise is the most fit; blue indicates that maximum volume module noise (10%) is the most fit; and green indicates that an intermediate level of volume module noise (5%) is the most fit.

**Fig 7.**
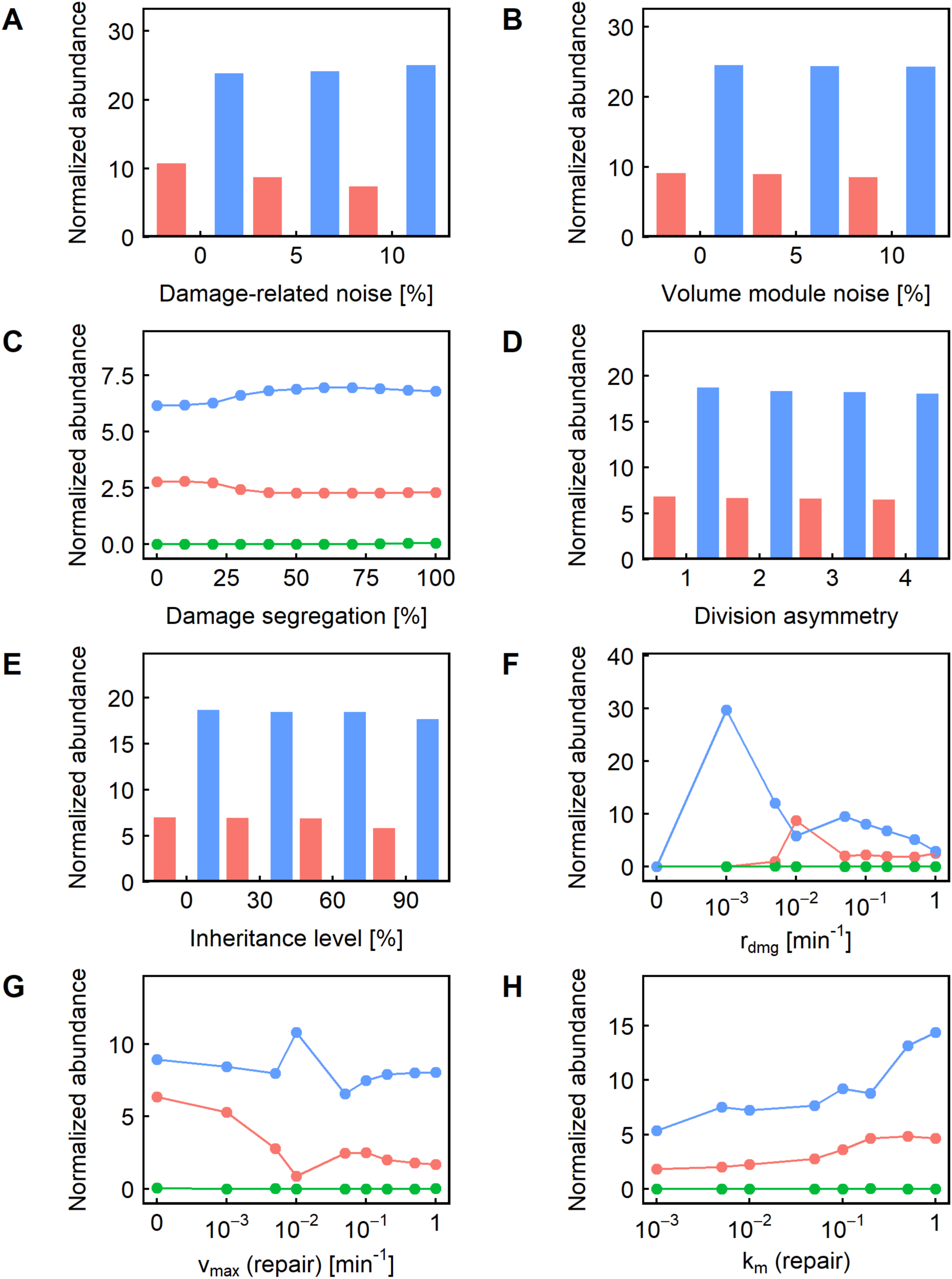
**A-H.** Relative representation of the other parameters in the cases where changing the nonlinearity of the effect of damage on volume growth rate caused a significant (>5%) fitness difference. In each panel, the color indicates the level of nonlinearity resulting in maximum fitness: red indicates that no nonlinearity (*α* = 1) is the most fit; blue indicates that maximum nonlinearity (*α* = 4) is the most fit; and green indicates that an intermediate level of nonlinearity is the most fit.

#### Effect of damage segregation on fitness

Somewhat surprisingly, we found that changes in damage segregation affected fitness in only 7% of the parameter value combinations used, compared to 6% for division asymmetry, and 7% for inheritance level and damage-related noise (Fig. 2B). To gain additional insights into what other parameters may interact with damage segregation to produce a fitness effect, we next examined those cases in which damage segregation could cause a significant change in population fitness. We found that most of the cases where damage segregation produced a fitness effect were seen when the damage rate was low and the repair rate was even lower (Fig. 3E, 3G), meaning that dilution is the primary method of damage reduction instead of active damage repair, giving significantly more prominence to the ability to sequester damage in the mother compartment. Another interesting observation related to these cases of parameter values is that higher values of *α* caused damage segregation to behave more like a double-edged sword: when *α* = 4, there were as many cases when damage segregation reduced fitness as when it improved fitness (Fig. 3F; compare red vs blue). This can be explained by the fact that the ultrasensitivity of fitness to damage level caused by the high nonlinearity diminishes the impact of damage segregation on fitness when the damage level is low but amplifies the effect when a threshold is crossed – in either direction. Once the damage rate becomes higher, however, damage segregation becomes more of a double-edged sword, causing fitness decreases about as often as it causes fitness increases. As a modular validation of the modeling approach we took in this study, consistent with previous reports [32], we also found that the highest damage rate (1 min^-1^) means that segregation is more likely to be beneficial compared to lower damage rates (0.1 or 0.2 min^-1^) (Fig. 3E).

#### Effect of division asymmetry on fitness

We next examined the effect of introducing division asymmetry on population doubling time or fitness. Introduction of division asymmetry caused significant changes (>5%) in doubling time in only 6% of the parameter value combinations used, making it the parameter the second least likely to alter the fitness outcome (Fig. 2B). Unlike damage segregation, the effects of division asymmetry on fitness are more likely to be double-edged: in 40% of the cases where a change in the division asymmetry parameter significantly altered fitness, the highest fitness was seen when there was no division asymmetry (Fig. 2B, red color). Like damage segregation, this behavior was mostly seen when the damage rate was low and the repair rate was even lower (Fig. 4E, 4G), corresponding to situations where dilution is the primary means of reducing damage levels. Moreover, such detrimental effects from division asymmetry were only seen when some damage segregation was present (Fig. 4C, red line). We interpret this result as follows. The smaller daughter cell usually takes longer to reach the volume threshold before it is ready to divide for the first time, which drags down the volume growth (and therefore dilution) rate at the population level. This effect is more pronounced when the damage segregation mechanism enriched the larger mother compartment with damage, slowing the growth of the mother compartment and dilution of damage. At higher rates of damage accumulation and repair, division asymmetry also becomes predominantly beneficial (Fig. 4E, 4G, blue line).

#### Effect of noise on fitness

Just as in the case for clonal senescence, we found that in most cases, higher noise levels had a beneficial effect on population fitness (Figs. 5 and 6, blue); this was particularly pronounced when the inheritance level was high (Figs. 5D, 6D). We interpret this as due to the high inheritance level permitting the propagation of “good” sets of parameter values selected by chance (and is more likely to be chosen if the noise value is high).

#### Summary

Overall, we found that model parameters directly affecting the accumulation of damage and its effect on the growth rate are the most likely to affect fitness. The introduction of damage segregation or division asymmetry, on the other hand, had a significant effect on population fitness in only a small fraction of the cases and in a context-dependent manner; however, when there was an effect, it was far more likely to be beneficial than detrimental. The fitness impact of asymmetric partitioning of cell volume was similarly context-dependent: when dilution was the predominant mechanism for damage removal, it was more likely to be detrimental, whereas if active damage repair was predominant, it was more likely to be beneficial.

## DISCUSSION

In this study, we comprehensively examine the effects of age-associated damage accumulation and removal on the phenotypes of clonal senescence and population fitness. Contrary to the results from a previous study which were based on a fully deterministic model with fixed parameter values [16], we found that neither damage segregation nor division asymmetry played a major role in the avoidance of clonal senescence once the natural diversity of the population is taken into account. Introduction of damage segregation eliminated clonal senescence only in a small fraction of borderline cases, while division asymmetry had an effect on clonal senescence in an even smaller fraction of cases; however, when there was an effect, it was more likely to cause clonal senescence than to eliminate it. While we acknowledge that the exact fraction will depend on the set of values and range chosen for the parameters, we believe that the relative value is still a good and useful indicator of the approximate importance and effect size of the parameters.

We note that our model differs from the model used in the previously published work in more ways than just the use of randomized coefficients. For instance, our model keeps track of single-cell volume explicitly, while the previous model used the protein count as a proxy for volume. Using protein count for cell volume inevitably led to some questionable assumptions where the partitioning of damaged proteins necessitated the inverse partitioning of undamaged proteins to maintain the protein count (and so the volume) of the daughter cell. Similarly, the sole mechanism for damaged protein to exert a detrimental effect in the previous model was by negative feedback on the production of new proteins, which required the amount of damaged proteins to be roughly comparable to that of intact proteins to have a meaningful effect, requiring likely unrealistic amounts of damage. Our model avoids these problems by using a more abstract “damage level” concept.

Uneven distribution of aging factors between daughter cells following cell division is a well-known phenomenon that has been observed in a variety of unicellular organisms, and asymmetric partitioning of volume has similarly also been observed in many unicellular organisms. In the present work, we comprehensively examine the fitness impact of these asymmetric damage and volume partitioning schemes and find that they, perhaps counterintuitively, have minimal fitness impact most of the time, as long as the natural diversity of the population is taken into account. When the repair rate was low and dilution was the predominant form of damage elimination, we found damage segregation to be more likely to be beneficial for fitness but division asymmetry to be generally detrimental. On the other hand, when active damage repair was the predominant damage elimination mechanism operating with a high damage repair rate, division asymmetry was found to be generally beneficial for fitness, while damage segregation had a double-edged impact, becoming more beneficial when the damage accumulation rate was very high.

Overall, our results here indicate that parameters governing the accumulation and elimination of cellular damage are the most important determinants of population fitness. Even though asymmetric partitioning of either cell volume or age-associated damage might seem beneficial at first glance, neither mechanism actually eliminates any damage on the population level, and, as we show here, they are far from being consistently beneficial evolutionarily. Why, then, are these mechanisms seen in some real organisms? To start with, the fitness impact of both mechanisms is context-dependent; depending on the values of other parameters, representing conditions both intrinsic and extrinsic to the cell, introduction of damage segregation and/or division asymmetry may be either beneficial or detrimental. Thus, it is possible that the organisms exhibiting damage segregation and/or division asymmetry fall within the section of the parameter space where such mechanisms are beneficial rather than detrimental, at least for some portion of their lifespan. And even if this region of the parameter space is not a common occurrence, the importance of avoiding irreversible senescence when it is encountered may be sufficient to preserve the mechanism evolutionarily, similar to how obscure nutrient pathways are preserved due to their critical importance under certain growth conditions. Second, in many cases, introduction of these mechanisms resulted in minimal fitness impact, but even minimal levels of fitness impact can accumulate over time and drive evolutionary selection. Moreover, several natural forms of damage are resistant to active repair, and thus may fall within the region of the parameter space where damage segregation is beneficial. For instance, carbonylated proteins can form aggregates that are resistant to proteasome digestion [33,34], and ERCs are self-replicating, suggesting that their effective repair rate – accounting for such self-replication – is probably also low [15,35]. Finally, for asymmetric partitioning of damage in particular, it has been reported that some organisms like *S. pombe* and *E. coli* only exhibit this behavior during high levels of external stress [14,18,32,36], suggesting that they may actually have evolved mechanisms to activate or inactivate damage partitioning depending on the region of parameter space they are in, just like other stress response mechanisms that are only activated in the presence of stress and can be epigenetically inherited by daughter cells [37,38].

## DECLARATIONS

### ACKNOWLEDGEMENTS

We thank G. Elison and G. Urbonaite for comments and feedback on the manuscript, and K. Miller-Jensen, C. O’Hern, and Acar Lab members for their comments on the project in its various stages. The authors also acknowledge the use of the Open Science Grid resources.

### AUTHOR CONTRIBUTIONS

RS and MA designed the project, including designing the model, simulations and their analyses. RS implemented the model in C++ and performed the simulations and analyses. RS and MA interpreted the results, and prepared, read, and approved the manuscript.

### DATA and MATERIAL AVAILABILITY

The data and materials related to this work are available upon request.

### ETHICS APPROVAL and CONSENT to PARTICIPATE

Not applicable.

### CONSENT to PUBLISH

Not applicable.

