## Supplementary Information for "Stochastic modeling of aging cells reveals how damage accumulation, repair, and cell-division asymmetry affect clonal senescence and population fitness"

**Supplementary Figures S1-S4**

**Supplementary Tables S1-S2**

### SUPPLEMENTARY FIGURES

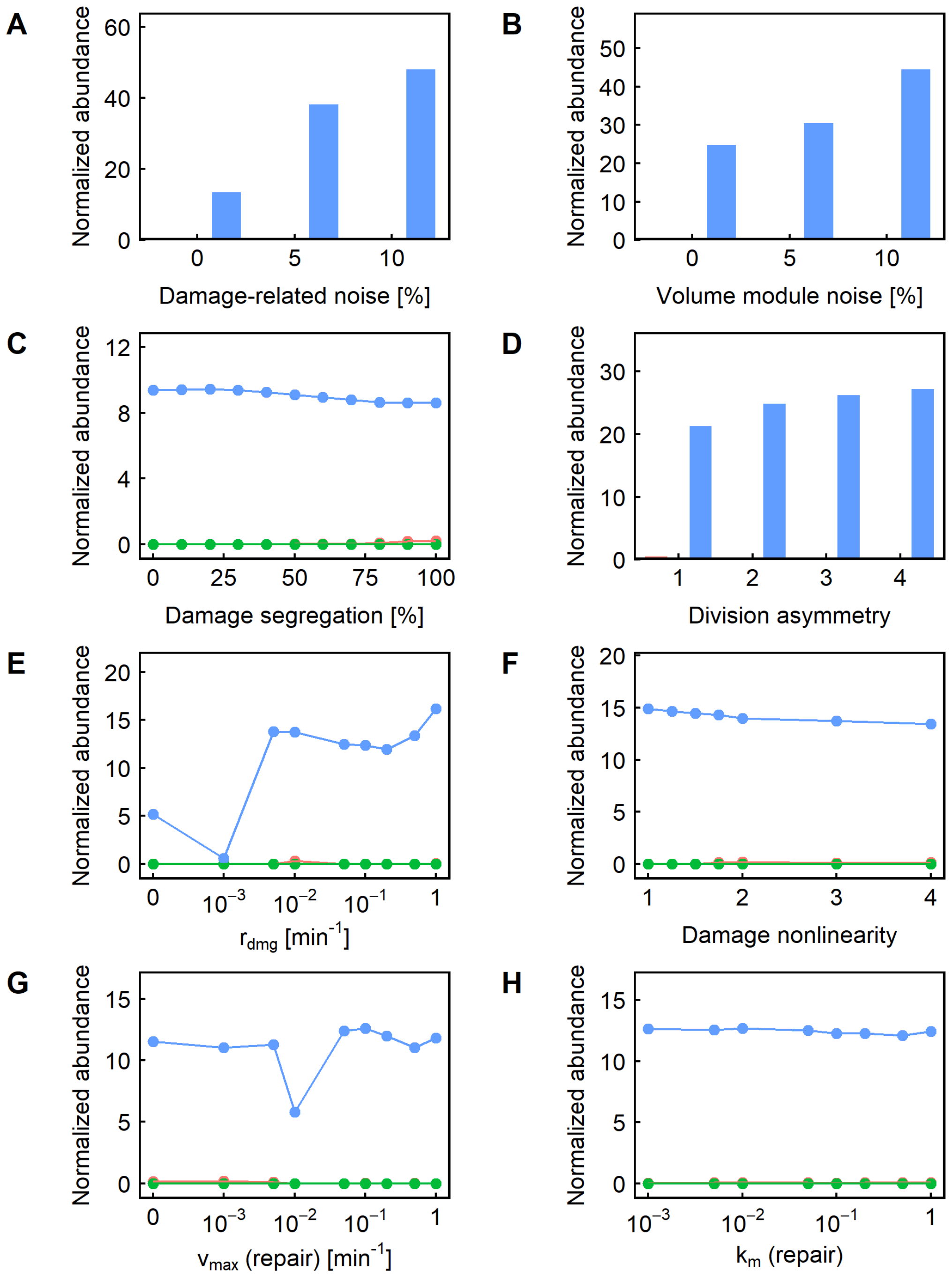


**Fig. S1. A-H.** Relative representation of the other parameters in the cases where changing the level of inheritance caused a significant (>5%) fitness difference. In each panel, the color indicates the level of inheritance resulting in maximum fitness: red indicates that no inheritance ($c$ = 0%) is the most fit; blue indicates that maximum inheritance ($c$ = 90%) is the most fit; and green indicates that an intermediate level of inheritance is the most fit.

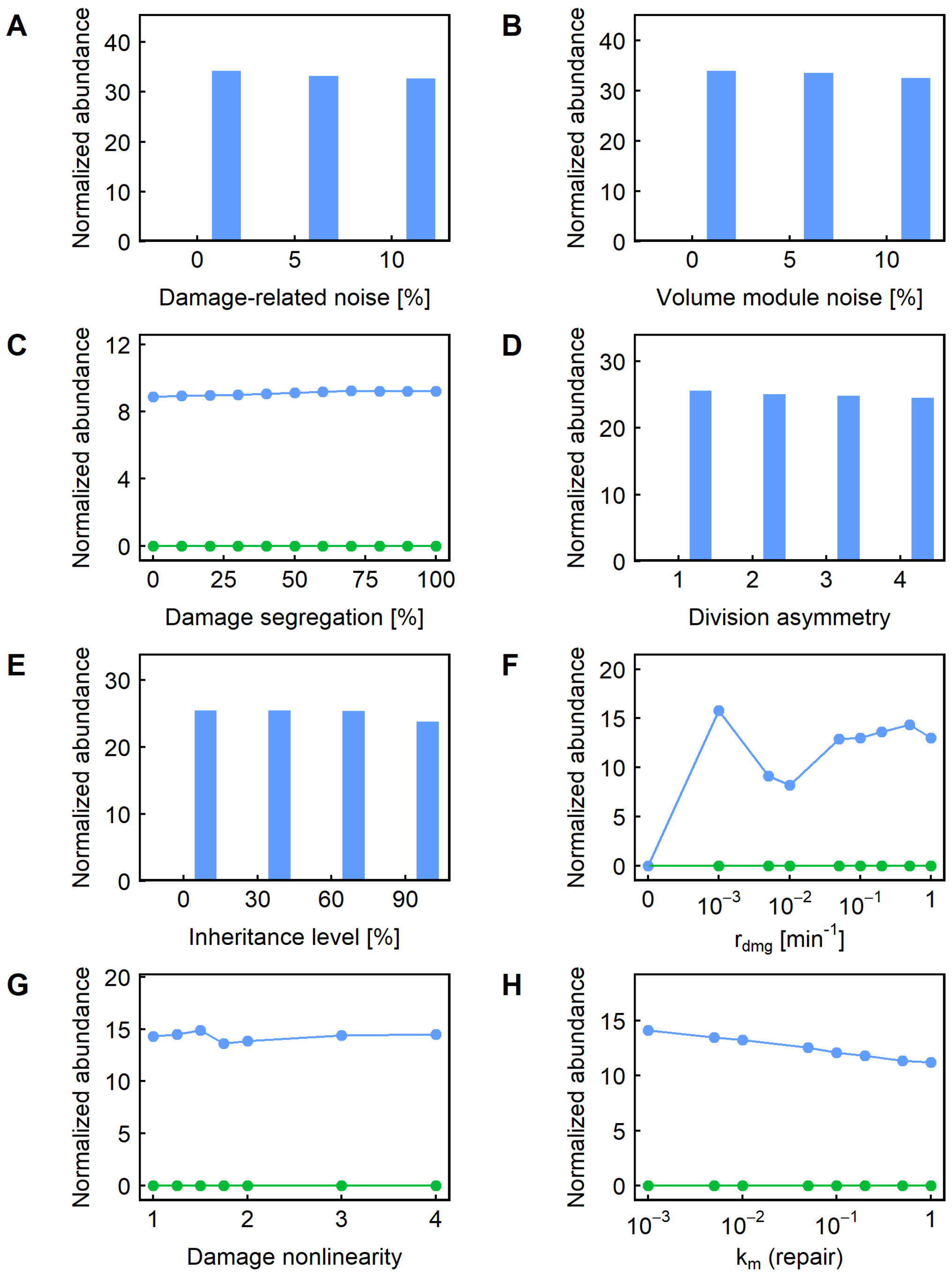


**Fig. S2. A-H.** Relative representation of the other parameters in the cases where changing the maximum repair rate caused a significant (>5%) fitness difference. In each panel, the color indicates the level of repair resulting in maximum fitness: red indicates that no repair is the most fit; blue indicates that maximum repair ($v_{max} = 1\,\text{min}^{-1}$) is the most fit; and green indicates that an intermediate rate of repair is the most fit.

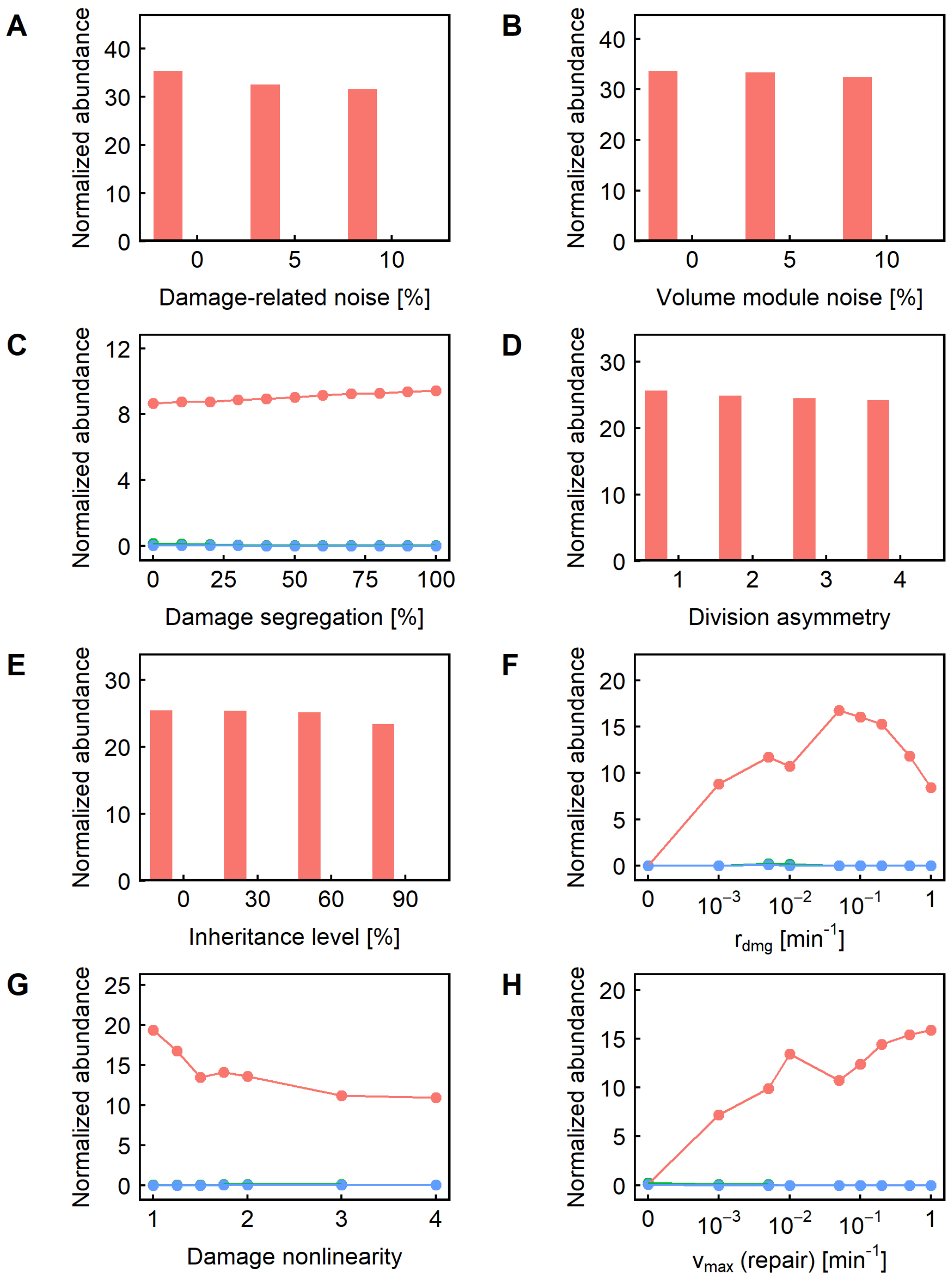


**Fig. S3. A-H.** Relative representation of the other parameters in the cases where changing the Michaelis constant $k_m$ for repair caused a significant (>5%) fitness difference. In each panel, the color indicates the value of $k_m$ resulting in maximum fitness: red indicates that the minimum value $k_m = 0.001$ is the most fit; blue indicates that maximum $k_m$ ($k_m = 1$) is the most fit; and green indicates that an intermediate value of $k_m$ is the most fit.

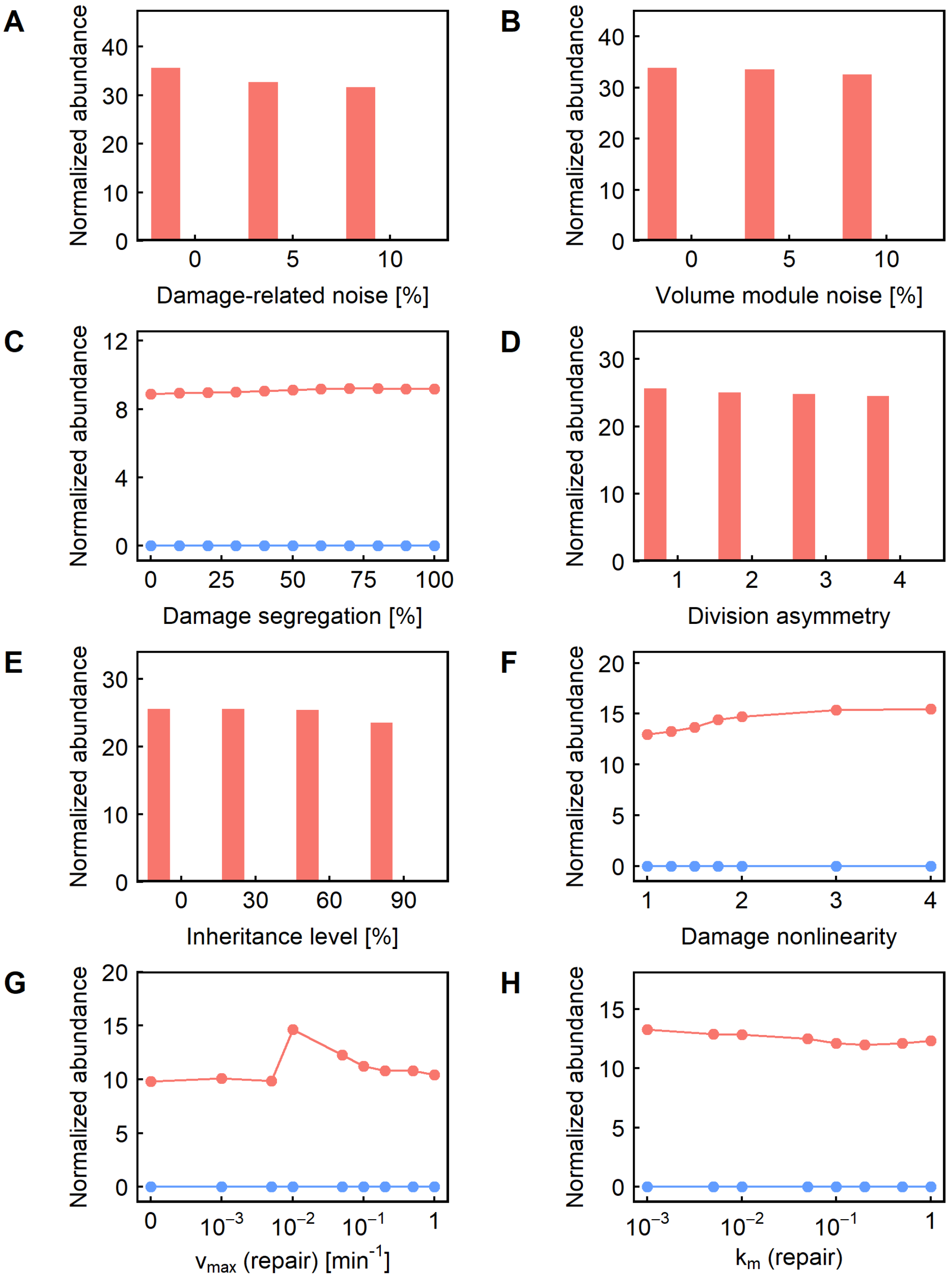


**Fig. S4. A-H.** Relative representation of the other parameters in the cases where changing the damage accumulation rate caused a significant (>5%) fitness difference. In each panel, the color indicates the damage accumulation rate resulting in maximum fitness: red indicates that no damage accumulation is the most fit; blue indicates that maximum damage accumulation ($r_{dmg} = 1\,\text{min}^{-1}$) is the most fit; and green indicates that an intermediate rate of damage accumulation is the most fit.

### SUPPLEMENTARY TABLES

| Parameter | Meaning | Value | Unit |
|---|---|---|---|
| $r_{growth}$ | Volume growth rate constant | 0.0072 | $min^{-1}$ |
| $V_i$ | Initial volume | 50 | fL |
| $V_{crit}$ | Critical volume (as function of generation *g*) | 106.7 + 4.628*g* | fL |

**Table S1. Parameters with fixed mean values.**

| Parameter | Meaning | Values | Unit |
|---|---|---|---|
| *c* | Level of inheritance of parameter values | 0, 30, 60, 90 | % |
| *R* | Mother/daughter volume ratio at division | 1, 2, 3, 4 | |
| $n_d$ | Damage-related noise | 0, 5, 10 | % |
| $n_v$ | Volume module noise | 0, 5, 10 | % |
| *s* | Level of damage segregation | 0, 10, 20, 30, 40, 50, 60, 70, 80, 90, 100 | % |
| $v_{max}$ | Maximum rate of repair | 0, 0.001, 0.005, 0.01, 0.05, 0.1, 0.2, 0.5, 1 | $min^{-1}$ |
| $k_m$ | Michelis constant for repair | 0.001, 0.005, 0.01, 0.05, 0.1, 0.2, 0.5, 1 | |
| $r_{dmg}$ | Rate of damage accumulation | 0, 0.001, 0.005, 0.01, 0.05, 0.1, 0.2, 0.5, 1 | $min^{-1}$ |
| *α* | Nonlinearity of the effect of damage on growth rate | 1, 1.25, 1.5, 1.75, 2, 3, 4 | |

**Table S2. Parameters with values selected from a grid of values.**
